# Thin Filament Interaction and Ca^2+^-Desensitization Effect of the C-Terminal End Peptide of Cardiac Troponin T Where Loss of Function Mutations Cause Hypertrophic Cardiomyopathy

**DOI:** 10.1101/2025.07.05.663331

**Authors:** Qiaobin Li, Han-Zhong Feng, J.-P. Jin

## Abstract

**Background:** Troponin T (TnT) is the tropomyosin (Tm)-binding subunit of troponin with a central role in regulating cardiac muscle contractility. The recently identified Tm-binding site 3 in the highly conserved C-terminal end segment of TnT has a troponin I (TnI)-like inhibitory function. Conformational modulation by proteolytic removal of the N-terminal variable region of cardiac TnT (cTnT-ND) in adaptation to inotropy-afterload mismatch enhances the function of Tm-binding site 3 to adjust ventricular contractile kinetics and sustain stroke volume. Mutations in this segment of cTnT cause overactivation of cardiac muscle and hypertrophic cardiomyopathy (HCM).

**Methods:** The C-terminal end 14 amino acid peptide of cTnT (cTnT-C14) was analyzed for interactions with Tm, F-actin, and F-actin–Tm thin filament using localized surface plasmon resonance (LSPR). Wild-type and HCM mutant cTnT-C14 peptides were used to treat skinned cardiac muscle strips from wild-type and cTnT-ND transgenic mice to assess the effect on Ca²⁺- activation of contraction.

**Results:** cTnT-C14 peptide showed saturable binding to Tm and F-actin in LSPR with physiological affinity, which is significantly impaired by HCM mutation R278C, K280N or R286C. Treatment of wild-type mouse cardiac muscle strips with cTnT-C14 peptide produced a Ca^2+^- desensitization effect on myofilament activation, which was not seen with cTnT-ND mouse cardiac muscle strips implicating a prior utilization of the same mechanism. The HCM mutant peptides lost this function.

**Conclusions:** The findings demonstrate the function of the cTnT C-terminal end segment underlying the pathophysiology of the HCM mutations. Its preserved functionality in the form of free peptide presents a drug candidate for the treatment of heart failure.

## Introduction

Cardiac muscle contraction and relaxation are essential for the pumping function of the heart. The contractile apparatus of vertebrate striated muscles is regulated by Ca^2+^ via the troponin complex in the sarcomeric thin filament. The troponin complex consists of three protein subunits: The Ca^2+^-binding subunit troponin C (TnC), the inhibitory subunit troponin I (TnI), and the tropomyosin (Tm)-binding subunit troponin T (TnT)^1^. During the activation of muscle contraction, cytosolic Ca^2+^ rises to bind TnC and induces a series of conformational changes in troponin and Tm, which in turn allows myosin heads to form strong cross bridges with the actin filament to activate myosin ATPase and a conformational power stroke that forces the sliding of thin filament toward the center of sarcomere to contract the muscle^2^.

TnT is at a central position in the thin filament regulation of skeletal and cardiac muscle contraction and relaxation. Extensive structure-function relationship studies have demonstrated that TnT binds TnI, TnC, Tm and F-actin^3^. Earlier studies have characterized two Tm-binding sites in the middle region and the beginning of the C-terminal region of TnT^4–6^. A very recent study further established a third Tm-binding site in the C-terminal end segment of TnT encoded by the last exon of vertebrate TnT genes^7,8^.

This C-terminal end segment of TnT has a highly conserved amino acid sequence among the cardiac, fast and slow skeletal muscle TnT isoforms and across vertebrate species^3^. A striking feature is that the 14 amino acids C-terminal end segment of cardiac TnT encoded by exon 17 of the *TNNT2* gene has identical sequence in all mammalian species (from platypus to human)^7,8^, indicating a stringently selected structure-function relationship. Consistently, the 14 amino acids C-terminal end segment of cardiac TnT is required for full inhibition of myofilament and substitution of the positively charged Arg and Lys residues in this segment with Ala resulted in increased activation of contraction^9^. Further demonstrating its functional importance, mutations within this segment, such as R278C, K280N or R286C, cause hypertrophic cardiomyopathy (HCM)^10–12^.

While TnC, the Ca^2+^-receptor subunit of troponin, belongs to the calmodulin family of Ca^2+^- binding proteins^13^, TnT and TnI, the two signal transmitting subunits of troponin, have evolved from duplication of a TnI-like ancestral gene^14^. Although TnT and TnI have significantly diverged in structures with specialized functions, removal of the evolutionarily added N-terminal variable region of TnT and its modulatory effect reconfigures the molecular conformation of the TnT backbone and brings back a TnI-like ancestral epitope structure^14^ detectable by an anti-TnI monoclonal antibody (mAb) that was raised against the C-terminal end segment of TnI^15^. The C-terminal end segment of cardiac TnI is critical to the inhibitory function and cardiac muscle relaxation^16^. The mAb TnI-1 recognized C-terminal peptide of cardiac TnI has been shown to bind Tm^17,18^. When a restrictive proteolytic deletion of the N-terminal variable region of cardiac TnT occurs in adaption to acute myocardial ischemia with contractility-afterload mismatch, the TnI C-terminal segment-like ancestral molecular conformation brought back in cardiac TnT produces a physiological reduction of the systolic velocity of cardiac muscle to elongate the left ventricular rapid ejection time and sustain stroke volume against afterload^19^. Pepride mapping studies have localized this TnI-like inhibitory Tm-binding activity to the 14 amino acids C-terminal end segment of TnT^7,8^.

To investigate the function of the C-terminal end segment of TnT as an inhibitory modulator of contractile kinetics of cardiac muscle, here we characterized the functionality of isolated C-terminal 14 amino acids peptide of cardiac TnT (cTnT-C14). Localized surface plasmon resonance (LSPR) spectroscopy determined the kinetics of the interactions of cTnT-C14 peptide with Tm and F-actin-Tm composite thin filaments. Its functional effect on Ca^2+^-activation of force production was determined in skinned cardiac muscle strips. The results provide new insights into the physiological function of the TnT C-terminal Tm-binding site 3 and the potential application of cTnT-C14 peptide as a novel reagent to adjust the kinetics of cardiac muscle contraction for the treatment of heart failure.

## 2. Materials and Methods

### 2.1. Amino Acid Sequence Analysis

TnT amino acid sequences were retrieved from NCBI GenBank database. The molecular mass and isoelectric point of peptides and their predicted charge at pH 7.0 were calculated using DNAStar software.

### 2.2. Synthetic Peptides

Wild-type (WT) human cTnT-C14 peptide and derivatives containing HCM mutation R278C, R286C or K280N were commercially synthesized by Peptide 2.0, Inc (Chantilly, VA) at >95% purity with HPLC mass spectrometry certification of the anticipated molecular mass. High concentration stocks of the peptides were made by dissolving them in a buffer containing 0.1 M KCl, 3 mM MgCl2, 10 mM imidazole-HCl, pH 7.0 and stored at -20°C until use in experiments. A non-Tm-binding and non-actin-binding peptide of the same length (E16) corresponding to the exon 16-encoded segment of human cardiac TnT was synthesized for use as a control.

### 2.3. Preparation of Tropomyosin and F-actin

Acetone powder of rabbit cardiac and back muscles was prepared from frozen tissue stocks. After minced using a precooled stainless-steel grinder and washed in cold water with stirring for 2-3 minutes and standing at 4°C for 20 minutes, the solubilized material was removed by squeezing out through two layers of cheesecloth. The tissue residue was sequentially washed at 4°C with 50% ethanol three times, 95% ethanol twice, and 100% acetone twice with stirring followed by filtered through cheesecloth. All equipment and reagents were precooled at 4°C. The resulting grayish-white acetone-washed residues were spread thinly over filter paper in a fume hood and air-dried at room temperature. The final dried acetone powder was stored at -20°C in aliquots until use.

As previously described^20^, α-tropomyosin was purified from cardiac muscle acetone powder by extraction with a buffer containing 1.0 M KCl, 0.5 mM dithiothreitol (DTT), 10 mM imidazole-HCl, pH 7.0, for 16 hours at room temperature with gentle stirring. The solubilized material was collected by squeezing out through two layers of cheesecloth, and the residue was re-extracted with the same buffer for 2 hours. The extracts were combined, adjusted to pH 4.6 using HCl, and stirred at 4°C for 30 minutes. The precipitate was collected by centrifugation at 6,000 × g at 4°C for 20 minutes and dissolved in the extraction buffer with stirring for 20 min. Any insoluble materials were removed by centrifugation at 6,000 × g at 4°C for 10 minutes. The isoelectric precipitation of tropomyosin at pH 4.6 and dissolution at pH 7.0 were repeated two more times. The final precipitate was dissolved in 0.5 mM DTT, 10 mM imidazole-HCl, pH 7.0. and precipitated on ice by slowly adding solid ammonium sulfate to 53% saturation at 0°C with stirring while maintaining pH at 7.0 with NaOH. After standing for 30 minutes, the precipitate was removed by centrifugation at 11,000 × g at 4°C for 30 minutes. More solid ammonium sulfate was added to the supernatant to 65% saturation at 0°C while keeping the pH at 7.0, and the precipitate was collected by centrifugation at 11,000 × g at 4°C for 30 minutes. The purified tropomyosin in the final precipitate was redissolved in deionized water containing 0.5 mM DTT and dialyzed against cold water containing 2 mM β-mercaptoethanol for three changes and lyophilized.

As described previously^21^, actin was extracted from rabbit back muscle acetone powder with a pre-cooled buffer containing 2 mM Tris-HCl, pH 8.0 containing 2 mM CaCl₂, 0.2 mM NaATP and 0.5 mM DTT with gentle stirring on ice for 30 minutes. The extraction mix was paper-filtered and centrifuged in a Beckman Coulter Optima XPN-100 Ultracentrifuge at 12,700 x g in a TYPE 50.2 Ti rotor at 4°C for 25 minutes to remove residual materials. KCl and MgCl₂ were gradually added to the supernatant to final concentrations of 50 mM and 2 mM, respectively, to initiate actin polymerization. This solution was gently stirred at room temperature for 1 hour. More KCl was added to reach a final concentration of 0.9 M followed by stirring at room temperature for 2 hours to separate tropomyosin from F-actin. The mixture was then ultracentrifuged as above at 244,000 x g at 4°C for 3 hours to pellet polymerized F-actin. The F-actin pellet was completely dissolved in a minimal volume of the extraction buffer and dialyzed against the same buffer at 4°C with twice changes to depolymerize F-actin. The solution of depolymerized actin was ultracentrifuged as above at 244,000 x g at 4°C for 2 hours to remove any insoluble materials. The clarified supernatant containing purified G-actin was repolymerized by adding KCl and MgCl₂ to 50 mM and 2 mM, respectively. The final product of F-actin was pelleted by ultracentrifugation as above at 244,000 x g at 4°C for 3 hours, resuspended in a small volume of the polymerization buffer and stored at 4°C for use within 7-10 days.

The concentration of rabbit α-tropomyosin and F-actin stocks was determined with UV absorbance at 280 nm based on their amino acid sequences.

### 2.4. SDS-Polyacrylamide Gel Electrophoresis (SDS-PAGE)

To assess the purity of Tm and actin prepared for the present study, SDS-PAGE was carried out as described previously^21,8^. The resolving gel is 14% with an acrylamide/bis-acrylamide ratio of 180:1. The protein samples were prepared in an SDS-gel sample buffer containing 2% SDS and 1% β-mercaptoethanol. The samples were heated at 80°C for 5 minutes and clarified by centrifugation in a microcentrifuge at top speed (20,000 x g) at room temperature for 5 min before loading to the gel. Electrophoresis was run at a constant current of 22 mA per 0.75 mm thick BioRad mini-gel. The gel was then stained with Coomassie Brilliant Blue R-250, distained, and scanned at 600 dpi for image documentation.

### 2.5. Localized Surface Plasmon Resonance (LSPR) Assay

LSPR spectroscopy was carried out using a Nicoya Lifesciences LSPR system to study the interaction between WT or mutant cTnT-C14 peptides and Tm or Tm-actin composite thin filaments. Nicoya High Capacity Carboxyl LSPR sensor chip was pre-coated with carboxyl-modified gold nanoparticles. The LSPR system was first flushed with 80% isopropanol at the maximum flow rate of 150 µL/min to remove any air bubbles. To activate the carboxyl groups on the gold nanoparticle surface before the coating of Tm or F-actin, 1-ethyl-3-(3- dimethylaminopropyl) carbodiimide (EDC) and N-hydroxysuccinimide (NHS) were dissolved separately in deionized water both at the concentration of 0.1 M. The EDC/NHS activation solution was made freshly by 1:1 mixing of the stocks immediately prior to use to ensure optimal activation efficiency. The EDC/NHS activation solution was injected at a flow rate of 20 µL/min for 4 minutes. The protein to be studied (Tm or F-actin) was immobilized on the chip surface by flowing the protein solution at 20 µL/min for 4 minutes for dehydration condensation of carboxyl groups on the chip with the amino groups on the protein.

For cTnT-C14 peptide-Tm binding assays, purified Tm was dissolved at 250 µg/mL in 10 mM sodium acetate, pH 4.0. It is important to have the pH at least 0.5 units below the isoelectric point of Tm (4.6) to ensure its positive overall charge. After Tm immobilization and equilibration in LSPR assay buffer consisting of 0.1 M KCl, 3 mM MgCl2 and 10 mM imidazole-HCl, pH 7.0, 1 M ethanolamine was injected at 20 µL/min for 4 minutes to block the unreacted carboxyl groups on the surface of gold particle on the chip. cTnT-C14 or control peptide diluted at 2 μM in the LSPR assay buffer was flowed over the chip at a rate of 20 µL/min for 4 minutes to record the course of association. The flow was then switched to the LSPR assay buffer at the same rate to elute the bound peptide and the course of dissociation was recorded until the signal level was back to baseline or reaching a plateau with no further decrease. The Tm-coated chip was regenerated between assays by washing with the LSPR assay buffer or with 1 M KCl and 3 mM MgCl2 at high flow rate for peptides with a strong binding to return the signal to baseline level.

For measuring the binding of cTnT-C14 to Tm-F-actin composite thin filament, F-actin was coated to the sensor chip same as that for Tm except for a protein concentration at 30 µg/mL. Tm in the LSPR buffer at 10 μM was applied to the F-actin-coated chip by flowing in at the rate of 20 µL/min for 4 minutes, allowing Tm to bind the immobilized F-actin at saturation. After a 2 minutes brief wash with the LSPR buffer at 20 µL/min to remove free Tm, cTnT-C14 peptide was injected at 2 mM with the flow rate of 20 µL/min for 4 minutes to measure its association to the immobilized Tm-F-actin filament. The flow was then switched to LSPR buffer and the dissociation of cTnT- C14 peptide and likely some Tm from the covalently immobilized F-actin was recorded. To regenerate the F-actin coated chip between assays, a 150 µL/min high-speed flow of the regeneration buffer containing 1 M KCl and 3 mM MgCl2 was applied to remove any residual Tm and TnT-C14 peptide.

The direct binding of cTnT-C14 peptides to chip-immobilized F-actin was measured similarly to the cTnT-C14-Tm binding assay.

Two of the HCM cTnT-C14 mutations, R278C and R286C, contain a cysteine residue which can form dimers via disulfide bonding. To prevent dimerization that could affect the binding properties for Tm or F-actin, 1 mM Tris(2-carboxyethyl)phosphine (TCEP) was added in the R278C and R286C stock solution at least 1 hour before testing to reduce disulfide bonds. After diluted Tm to 2 µM in the LSPR working solution the final TCEP concentration was 20 µM, which did not generate any background signal while ensuring any disulfide bonds were reduced.

### 2.6. Measurements of Ca^2+^-Activated Force of Skinned Cardiac Muscle Strips

The animal procedure was carried out using a protocol approved by the Animal Care Committees of University of Illinois at Chicago and were conducted in accordance with the National Institutes of Health Guide for the Care and Use of Laboratory Animals.

Left ventricular papillary muscles from 3-4 months old WT C57B/L6 mice or transgenic mice expressing N-terminal truncated cardiac TnT (cTnT-ND)^19^ were isolated immediately after euthanasia and flash frozen at their slack length. Using the method described previously^22^, 35 μm thick longitudinal cryosections were prepares and then longitudinally sliced into 120-150 μm wide cardiac muscle strips. Glycerol-permeabilized cardiac muscle strips were mounted between two aluminum T-clips and transferred to a chamber of a thermo-controlled stage (802D, Aurora Scientific) at 6-8°C in a relaxation buffer (BES 40 mM, EGTA 10 mM, MgCl2 6.86 mM, ATP 5.96 mM, DTT 1 mM, creatine phosphate 33 mM, creatine kinase 200U/mL, K-propionate 3.28 mM, pH 7.0, plus protease inhibitor cocktail). For contractility studies, the muscle strip was connected to a force transducer (403A, Aurora Scientific) and a length controller (322C, Aurora Scientific). The buffer was then switched to a skinning solution (relaxation buffer containing 1% Triton X-100) for 20 min. After a wash with relaxation buffer, the fully permeabilized muscle strip was placed in pCa 9.0 buffer and the sarcomere length was measured through a digital camera attached to the microscope and adjusted to 2.0 μm or 2.3 μm for mechanical measurements. After measuring the baseline calcium activated force was examined at pCa 6.5, 6.3, 6.0, 5.8, 5.5, 5.0, and 4.5 at 15°C, WT or HCM mutant HcTnT-C14 peptides or the control peptide was added at 20 μM and the force-pCa measurements repeated. The force development of the muscle strip at pCa 4.5 was then measured to verify the absence of any significant rundown effect.

### 2.7. Data Analysis

Statistical comparisons of LSPR-determined Ka, Kd, and KD results were done using unpaired Student’s t test. All values are presented as mean ± SD. The force-pCa curves were plotted and fitted using a Hill exponential equation (y = START + (END-START)*x^n / (k^n+x^n)) for data analysis and statistical analysis was performed using paired Student’s t-test. All values are presented as mean ± SE.

## 3. Results

To illustrate the foundation for investigating the functions of WT and HCM mutant cTnT- C14 peptides, the C-terminal end location of cTnT-C14 peptide in reference to the recently identified Tm-binding site 3 and other functional sites of TnT is shown in Fig. 1A. Fig. 1B shows an amino acid sequence alignment of cTnT-C14 segment of representative mammalian species and the striking feature that this segment is 100% conserved in all mammalian hearts, indicating a stringent evolutionary selection with functional importance. Physical properties of the TnT-C14 peptides were compared in Fig. 1B to demonstrate that the loss of a positive charge in the MHC mutants may be a common pathogenic factor. Supporting this notion, previous studies have shown that substituting the positively charged residues in the C-terminal end segment of TnT with neutral residue Ala resulted in impaired relaxation and overactivation of skeletal and cardiac muscles^9,23^.

**Figure 1.**
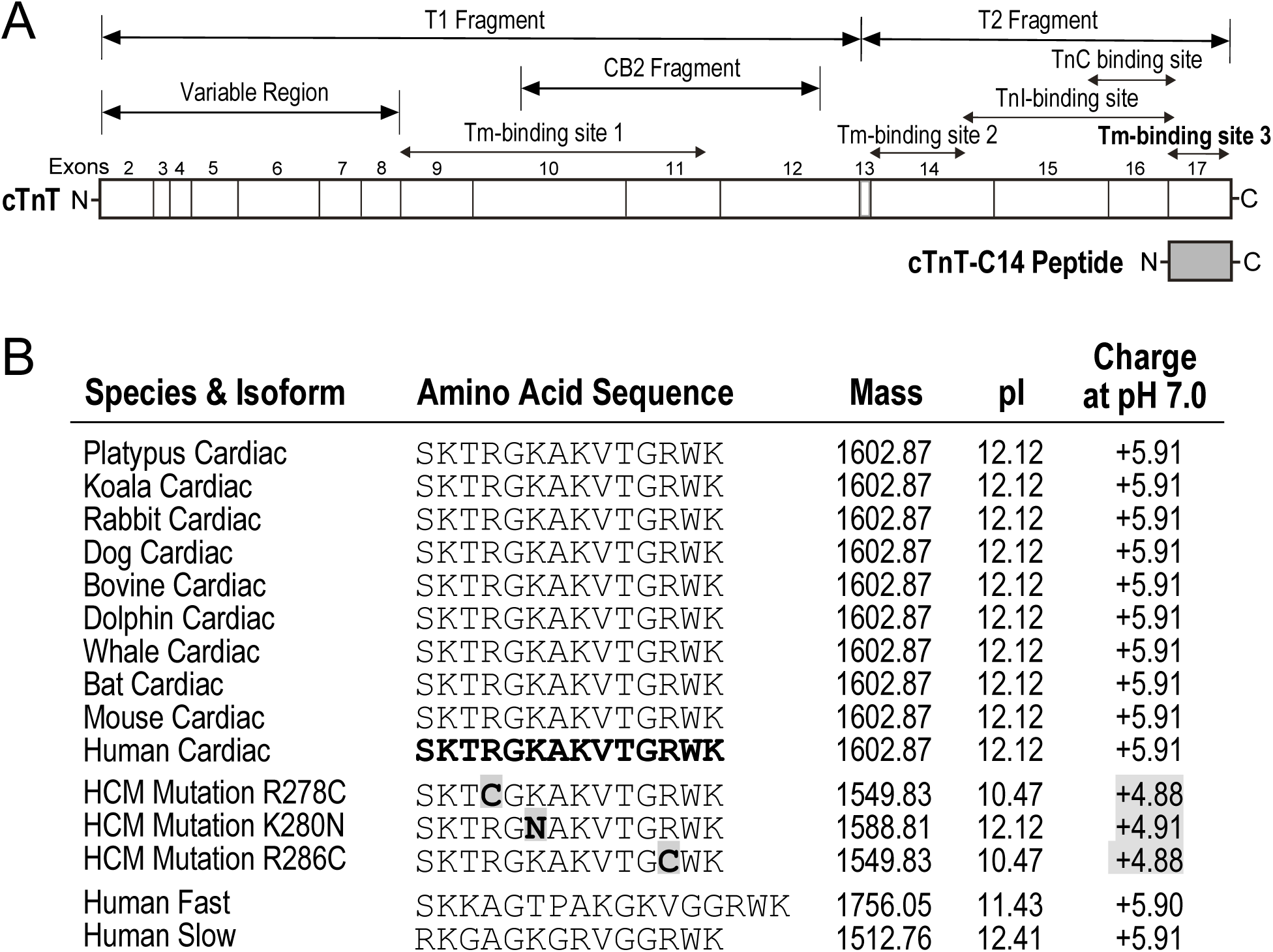
The C-terminal end segment of mammalian cardiac TnT in comparison with skeletal muscle isoforms and cardiomyopathy mutations. (A) Primary structure map of cardiac TnT is aligned with the cTnT-C14 peptide to show their structural relationships with the segments encoded by each exon and the binding sites for TnI, TnC and Tm in relation to the T1 and T2 chymotryptic and CB2 CNBr fragments indicted. (B) The amino acid sequence alignment of the C-terminal end segment of TnT along with the average isoelectric point (pI) and net charge at pH 7.0 showed that structure of the C-terminal end segment encoded by the last exon of TnT genes is highly conserved and has an identical amino acid sequence in all mammalian species from platypus to human. The C-terminal end segments of human fast and slow skeletal muscle TnT are aligned as examples to show the variation among muscle type isoforms. The three cTnT- C14 peptides containing hypertrophic cardiomyopathy (HCM) mutation R278C, K280N or R286C are compared to implicate that the loss of a positive charge may be a pathogenic factor shared by these mutations. The NCBI GenBank accession numbers of the TnT sequences analyzed are: Platypus cardiac, XP_028924875.1; koala cardiac, XP_020858612; rabbit cardiac, AAB51160.1; dog cardiac, AAG23715.1; bovine cardiac, NP_777196.1; dolphin cardiac, XP_059982389.1; whale cardiac, XP_057400554.1; bat cardiac, XP_066095192.1; mouse cardiac, BAB19881.1; human cardiac, AAK92231.1; human fast, NP_001354775.1; and human slow, NP_001119604.1.

The SDS-PAGE gels in Fig. 2 show that the purified Tm and F-actin used in the functional studies had no detectable contamination of other proteins.

**Figure 2.**
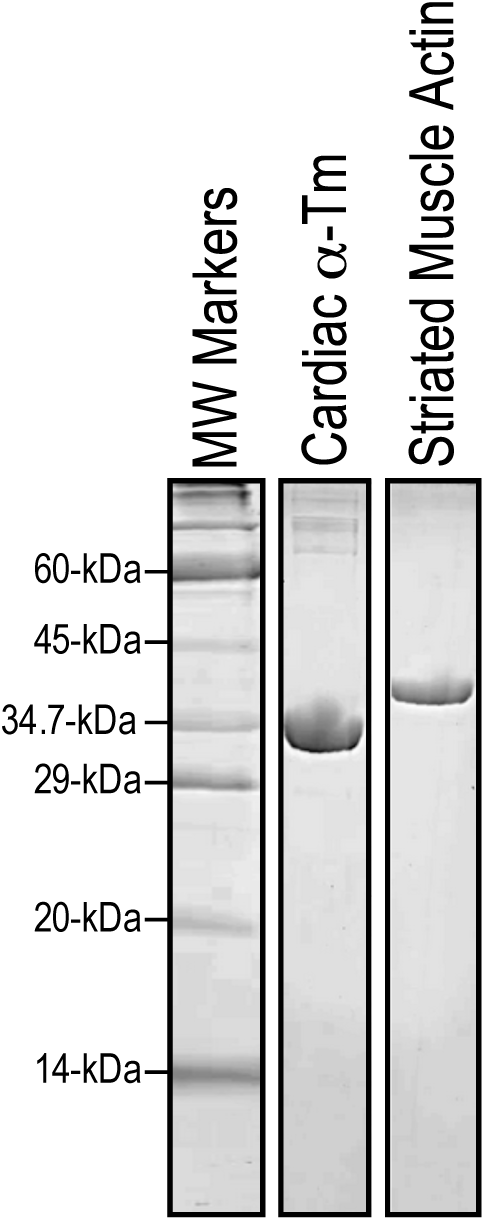
Purified cardiac α-tropomyosin and striated muscle actin used in the present study. The SDS-gel images show the tropomyosin and actin preparations of good purity and anticipated molecular weights (43-kDa and 32.5-kDa, respectively).

### 3.2. Tm-Binding Kinetics of cTnT-C14 Peptide and the Negative Impact of HCM Mutations

The representative LSPR traces in Fig. 3 show that WT human cTnT-C14 peptide at 2 μM concentration binds immobilized Tm with a rapid association rate to reach saturation. Once saturation was reached and the flow was switched to LSPR buffer, the dissociation of WT cTnT- C14 peptide occurred also rapidly. The physiologically relevant binding affinity with fast kinetics (Table 1) demonstrate that functionality of the Tm-binding site 3 in the C-terminal end segment of TnT is retained in the form of isolated free peptide. The high Tm-binding affinity of isolated cTnT- C14 peptide demonstrates an intrinsic conformational and functional state of the Tm-binding site 3 of TnT, which is known to be conformationally modulated by the N-terminal variable region (Fig. 1A)^7,8^. The fast association and dissociation kinetics are consistent with the function of Tm-binding site 3 in modulating contractility of cardiac muscle during the dynamic pumping cycle of the heart.

**Figure 3.**
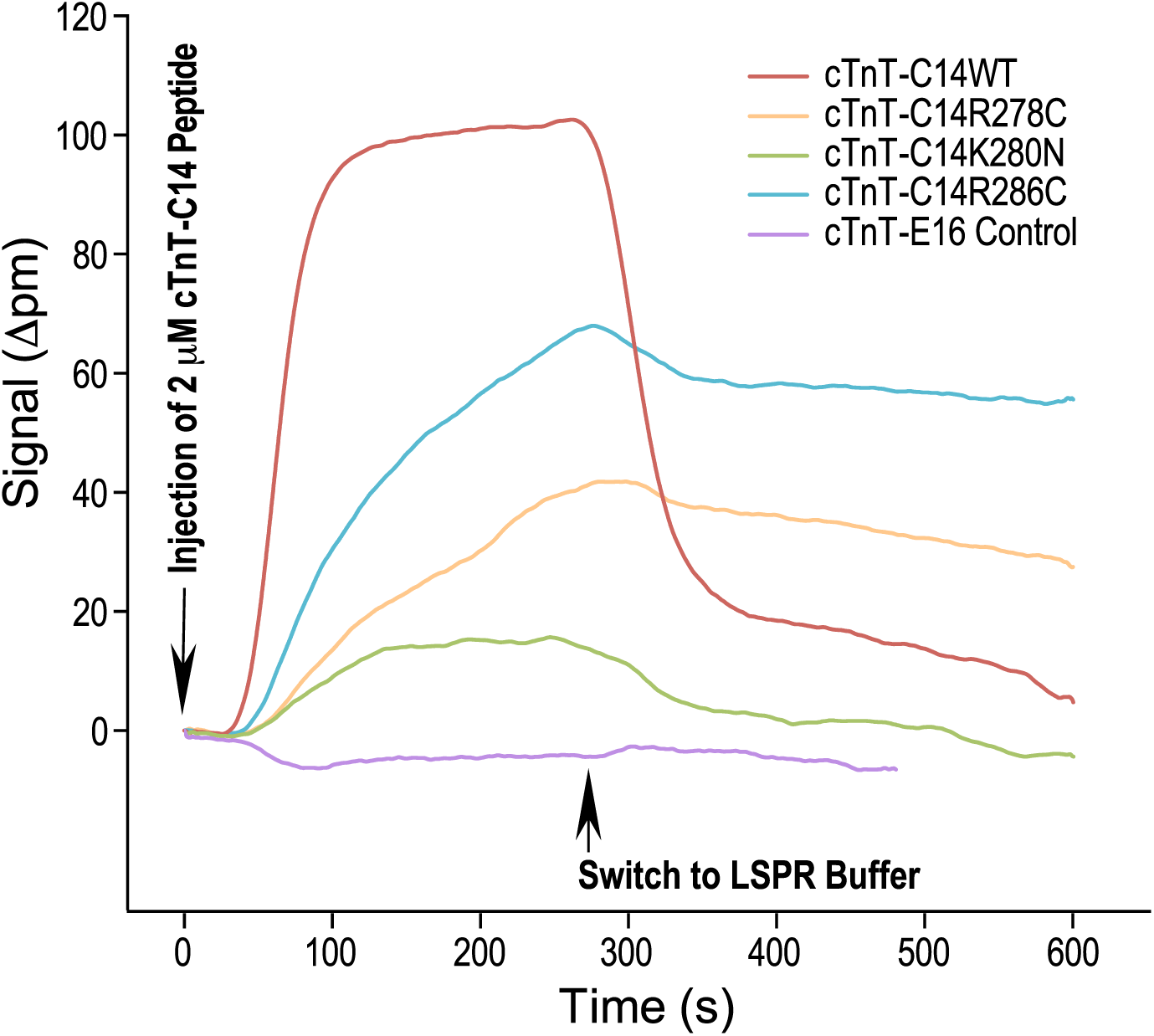
LSPR analysis of the binding of cTnT-C14 peptides and tropomyosin. WT human cTnT-C14 peptide or cTnT-C14 peptides containing HCM mutations R278C, K280N and R286C at 2 μM concentration was flow through an LSPR sensor chip with saturated coating of Tm. In comparison with the negative control peptide cTnT-E16, the representative LSPR traces from three independent experiments demonstrate that cTnTC-14WT produced a saturable binding to immobilized Tm with fast association and dissociation rates. In contrast, the three HCM mutant peptides all showed weakened Tm-binding with significantly decreased association rates. While HcTnT-C14K280N shows much diminished binding affinity for Tm, HcTnT-C14R278C and HcTnT-C14R286C show decreased maximum binding but notably slower dissociation rates. The results demonstrate that the C-terminal end segment of cardiac TnT functions as a Tm-binding site in the form of isolated free peptide at physiologically relevant affinity, which is significantly impaired by the HCM mutations.

**Table 1.**
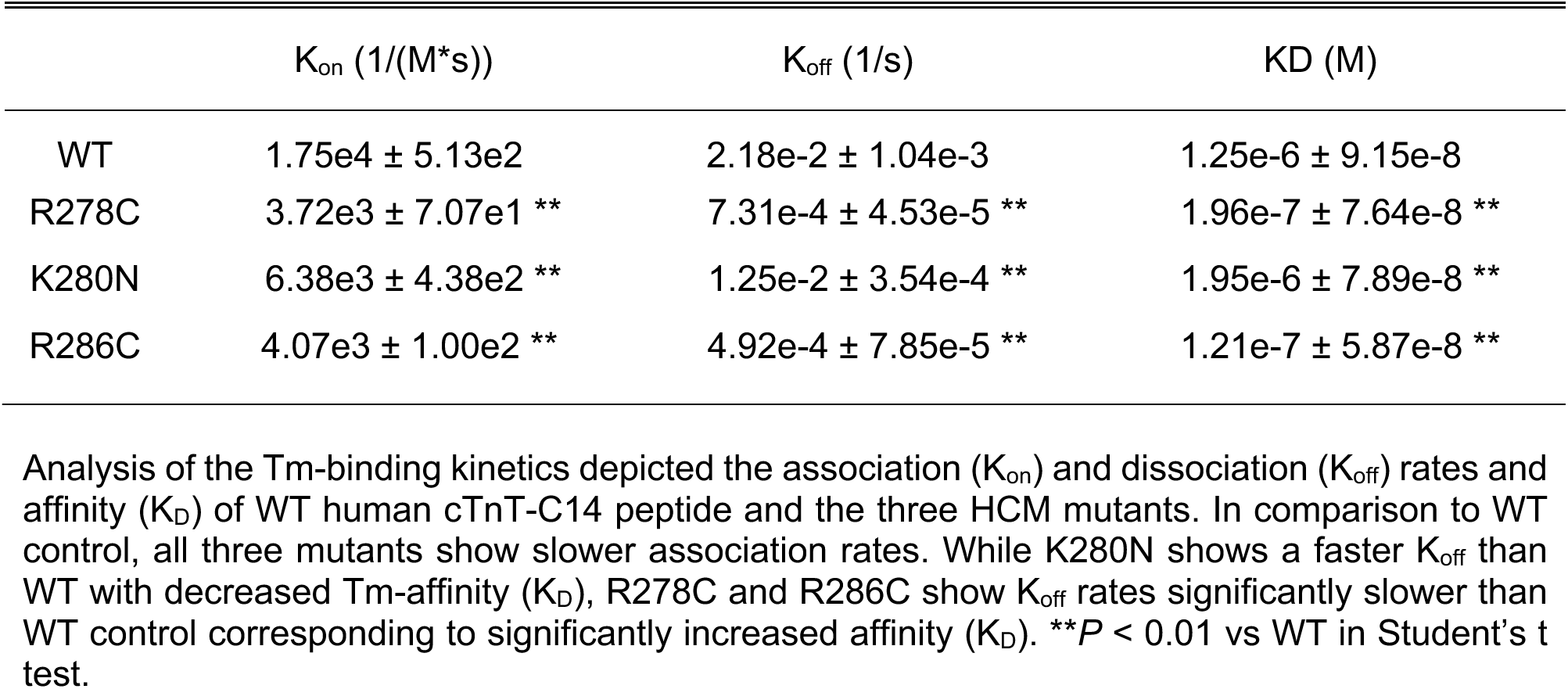
Association (Kon) and dissociation (Koff) rates and affinity (KD) of the binding of WT and HCM mutant human cTnT-C14 peptides to immobilized Tm in LSPR.

The results in Fig. 3 and Table 1 further show that human cTnT-C14 peptides containing one of the three single amino acid HCM mutations (R278C, K280N or R286C) all have significantly decreased binding to Tm. In comparison to WT cTnT-C14 under the same concentration, time parameters and experimental conditions, the maximum Tm-binding signal of the mutant peptides were significantly lower, most notably for the K280N mutant which dropped to nearly that of the negative control peptide.

Consistent with weakened Tm-binding affinity, the HCM mutant peptides all displayed significantly slower association rates (Kon). Interestingly, R278C and R286C also show slower rates of dissociation (higher Koff) (Fig. 3 and Table 1), indicating that once the mutant cTnT-C14 peptides bind to Tm, they form more stable complex than that of the WT peptide. The slower kinetics indicate weaker and less dynamic interactions with Tm, which may underlie the pathophysiological effects of the HCM mutations on damping the cardiac muscle contraction-relaxation cycle to decrease pumping efficiency.

### 3.2. Binding of cTnT-C14 Peptide to Tm-F-actin Composite Thin Filament and Impact of the HCM mutations

The LSPR results in Fig. 4 and Table 2 show that WT human cTnT-C14 peptide binds to Tm-F-actin composite filaments with high affinity, similar to its saturable binding to Tm alone with fast association and dissociation rates. Also similar to the binding to Tm alone, cTnT-C14 peptide containing the HCM mutant, especially K280N, exhibited significantly weakened binding to Tm-F-actin composite filament. The R278C and R286C mutants demonstrated slower association rates than that of WT control. The binding kinetics of cTnT-K280N could not be determined from the very weak binding curve. The dissociation trace signal represents a complex process including the dissociation of both cTnT-C14 peptide and Tm from the solid phase, which precludes specifically calculating the dissociation rate of the cTnT-C14 peptides from Tm-F-actin filament.

**Figure 4.**
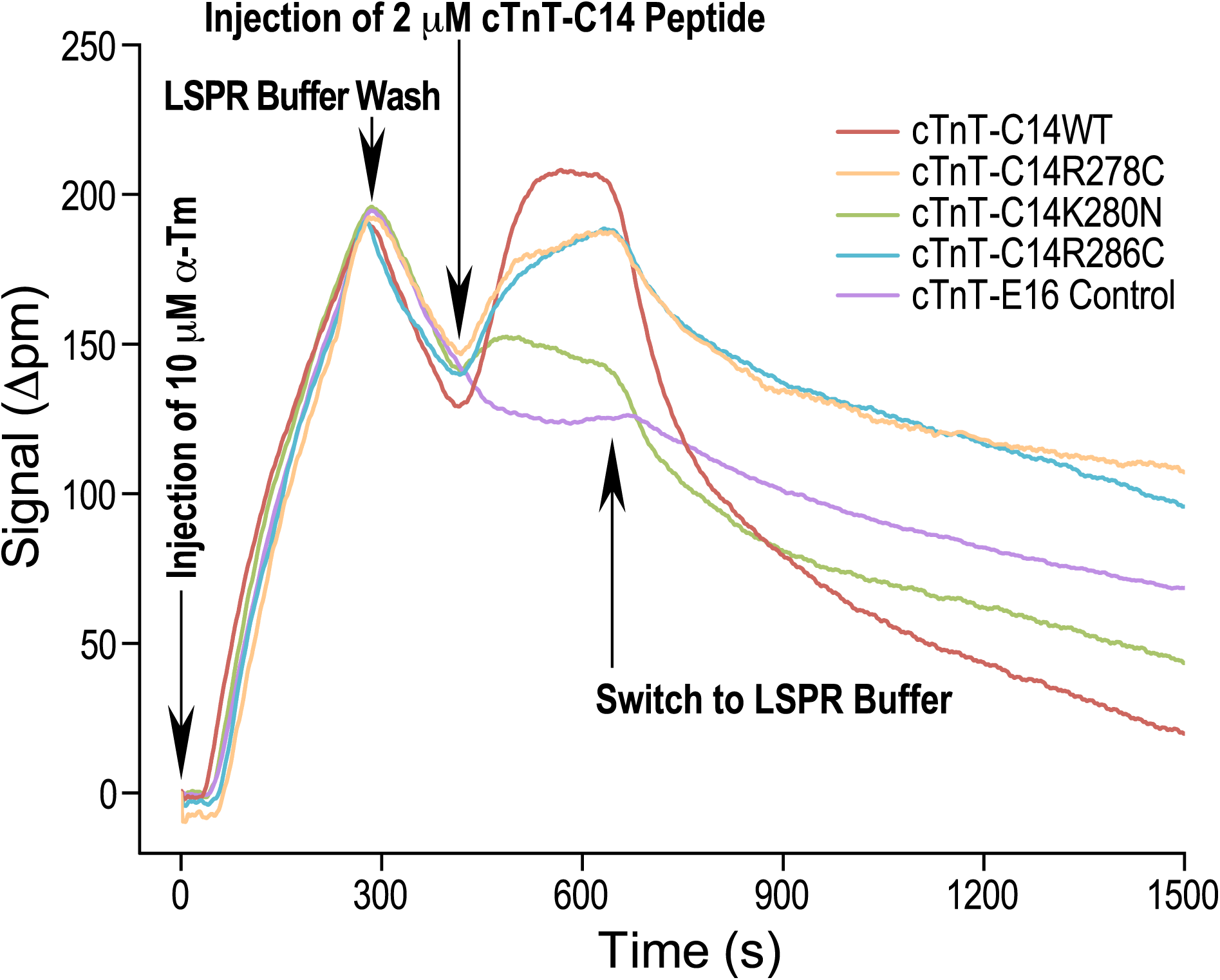
LSPR analysis of the binding of cTnT-C14 peptides to tropomyosin-actin composite filament. The sensor chip was coated with a low density of F-action (at the flow concentration of 30 μg/mL). After a saturating addition of Tm (at the flow concentration of 10 μM) to the immobilized F-actin and a brief wash to remove free Tm, WT or HCM mutant human cTnT- C14 peptides was injected at 2 μM concentration to measure the association to and dissociation from the Tm-F-actin filaments. In comparison with the negative control peptide cTnT-E16, the LSPR profile shows an initial peak representing the binding of Tm to F-actin followed by the binding peak of cTnT-C14 peptides. The representative LSPR traces from at least three independent experiments demonstrate that WT human cTnT-C14 peptide had rapid association and saturable binding to Tm-F-actin filament followed by a rapid dissociation, similar to its interaction to Tm alone (Fig. 3). Also similar to the binding patterns to Tm alone, the K280N mutant showed drastically diminished binding affinity to the Tm-F-actin filament while the R278C and R286C mutants showed decreased maximum binding with slowed dissociation rates. The results confirm that the C-terminal end peptide of cardiac TnT binds Tm in muscle thin filament with physiological relevance, which is impaired by the HCM mutations.

**Table 2.** Association (Kon) rate of WT and HCM mutant cTnT-C14 peptides to immobilized Tm-F-actin composite filament.

| | $K_{on}$ (1/(M*s)) |
| --- | --- |
| WT | $1.08e4 \pm 8.34e2$ |
| R278C | $7.73e3 \pm 4.95e1$ * |
| R286C | $6.77e3 \pm 7.23e2$ * |

Binding kinetic analysis of the association rate ( $K_{on}$ ) to Tm-F-actin composite thin filament showed slower binding of the R278C and R286C mutants than WT control. The binding kinetics of cTnT-K280N was not calculated because of the very weak binding curve that precluded a reliable fitting analysis. The dissociation rate of the cTnT-C14 peptides from Tm-F-actin filament could not be specifically calculated since the dissociation trace signal represents a complex process including the dissociation of both cTnT-C14 peptide and Tm from the solid phase. \* $P < 0.05$ vs WT in Student's t test.

These results further support the physiological relevance of the Tm-binding function of cTnT-C14 peptide in adjusting muscle contractility. The similar patterns of WT cTnT-C14 peptide interactions with Tm-F-actin filament and with Tm alone, and the similar effects of each the three HCM mutants, indicate that the binding to Tm is the primary functional mechanism of the cTnT- C14 peptide, which is predominant in the binding to Tm-F-actin composite thin filament. The changes in cTnT-C14 peptide’s thin filament-binding kinetic caused by the HCM mutations demonstrate a pathogenic and pathophysiological mechanism through impairing the Tm-binding site 3-based regulation of thin filament Ca^2+^ activation and deactivation and the kinetics of cardiac muscle contraction and relaxation.

### 3.3. cTnT-C14 Peptide Directly Binds F-actin

The C-terminal domain of TnT has previously been indicated with an activity in binding F-actin^24^. Knowing the cTnT-C14 peptides bind Tm-F-actin composite filament similarly to their binding patterns for Tm alone (Figs. 3 and 4) indicating the primary interaction with Tm, we further investigated their direct binding to F-actin in LSPR under the same conditions. The results in Fig. 5 and Table 3 show that WT cTnT-C14 peptide binds immobilized F-actin with a physiologically relevant affinity but weaker than that for Tm alone and did not reach saturation during the 300 s flow time. The F-actin-binding kinetics show a slightly slower association rate and a slightly fast dissociation rate (Table 3) than that to Tm (Table 1). The HCM mutations also decreased the binding of cTnT-C14 peptides to F-actin (Table 3) while the dissociation rates of R278C and R286C were slower than WT control (Table 3), similar to that with Tm (Table 1).

**Figure 5.**
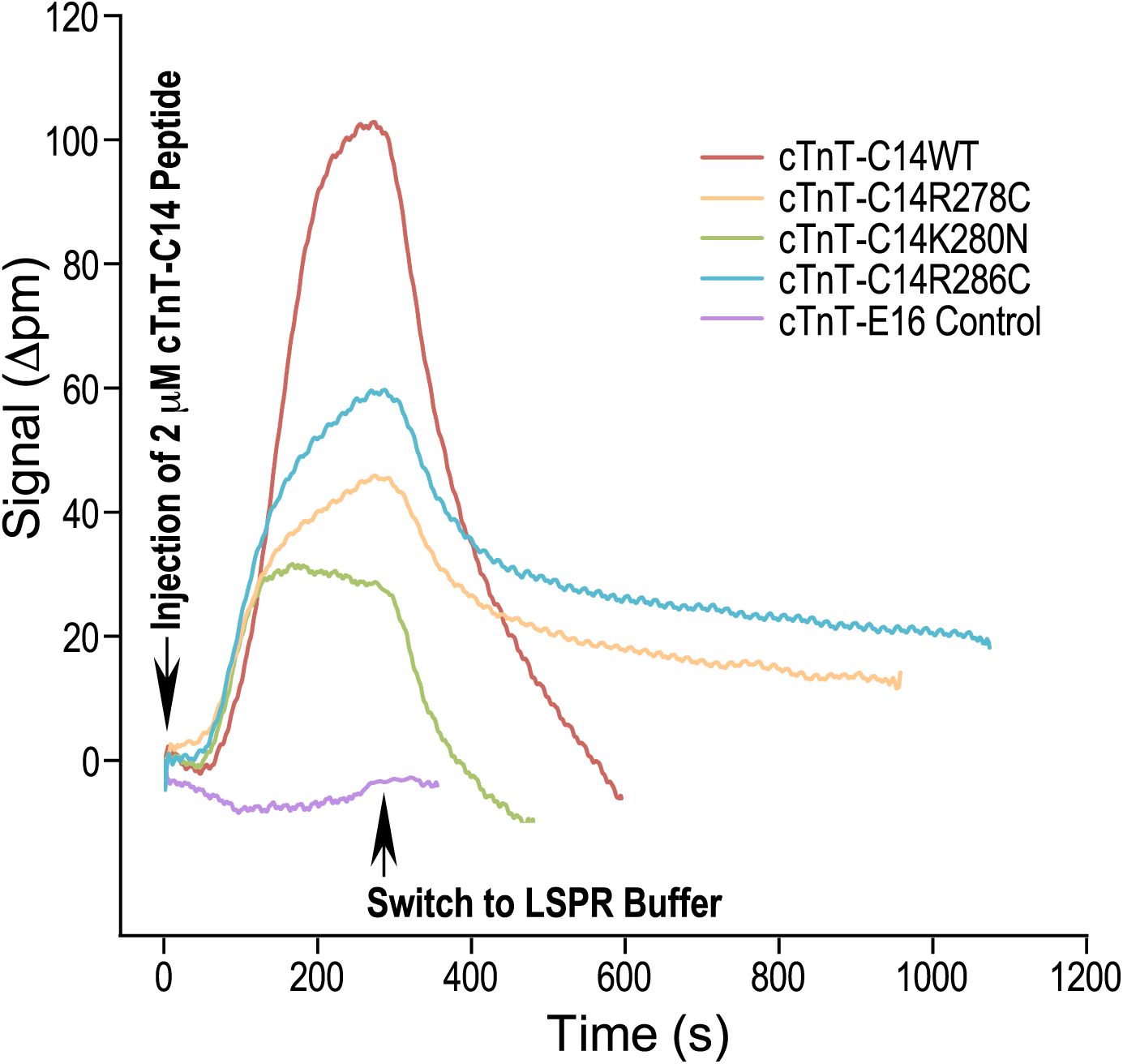
LSPR analysis of the direct binding of cTnT-C14 peptides to F-actin. LSPR sensor chip was coated with F-action (at the flow concentration of 30 μg/mL) to test the interactions of WT and mutant human cTnT-C14 peptides with F-actin. The representative LSPR traces show that cTnT-C14WT produces the strongest binding in contrast to the weaker bindings of the mutants. Distinct from the binding kinetics to Tm of Tm-F-actin thin filaments, the binding of cTnT- C14 WT was slower and did not reach a plateau in 300 s. The binding of cTnT-R278C and cTnT- R286C showed lower maximum binding to F-action with slower rates of dissociation similar to that from Tm alone (Fig. 3). The binding of cTnT-C14K280N to F-actin exhibits fast association and dissociation rates but a significantly lower level of maximum binding. The similarities and differences of WT and HCM mutant cTnT-C14 peptides’ binding kinetics with Tm and F-actin suggest that the interaction of TnT C-terminal end segment with Tm and actin are via different mechanism involving different amino acid residues.

**Table 3.** Association (Kon) and dissociation (Koff) rates and affinity (KD) of the direct binding of cTnT-C14 peptides to immobilized F-actin in LSPR.

| | $K_{on}$ (1/(M*s)) | $K_{off}$ (1/s) | KD (M) |
| --- | --- | --- | --- |
| WT | $8.02e3 \pm 9.90e1$ | $9.84e-3 \pm 5.98e-4$ | $1.41e-6 \pm 2.53e-7$ |
| R278C | $6.41e3 \pm 6.00e2$ | $4.86e-3 \pm 2.76e-4$ ** | $7.61e-7 \pm 4.46e-8$ * |
| K280N | $1.23e4 \pm 1.67e3$ ** | $1.50e-2 \pm 9.85e-4$ ** | $1.24e-6 \pm 2.32e-7$ |
| R286C | $6.54e3 \pm 4.31e2$ | $5.38e-3 \pm 2.91e-4$ ** | $1.08e-6 \pm 4.66e-7$ |

Analysis of F-actin-binding kinetics depicted the association ( $K_{on}$ ) and dissociation ( $K_{off}$ ) rates and affinity ( $K_D$ ) of WT human cTnT-C14 peptide and the three HCM mutants. In comparison to WT cTnT-C14, the K280N mutant showed detectably faster association and dissociation rates whereas the R278C and R286C mutants showed slower rates of dissociation than WT control. \* $P < 0.05$ ; \*\* $P < 0.01$ vs WT in Student's t test.

The results demonstrate that the C-terminal end segment of TnT is involved in the previously detected direct binding of TnT to F-actin^24^. The differences in the kinetics of Tm-binding and F-actin-binding, such as the different association rates but similar dissociation patterns, indicate distinct binding mechanisms and involvement of possibly different amino acid residues in the C-terminal end segment of TnT. This notion is supported by an observation that the negative impact of the K280N mutation on F-actin-binding (Fig. 5) is less than that on Tm-binding (Fig. 3), indicating distinguishable structural bases. Altogether, the similar patterns of cTnT-C14 peptides’ binding to Tm-F-actin thin filaments and Tm alone support that the functionality of the C-terminal end segment of TnT in regulating muscle contractility is primarily via its Tm-binding activity.

### 3.4. Effects of cTnT-C14 Free Peptide on the Contractility of Skinned Cardiac Muscle

Force-pCa studies showed that the addition of WT cTnT-C14 peptide to skinned WT mouse left ventricular papillary muscle strips decreased the Ca^2+^-sensitivity for activation of contraction (Fig. 6A). The treatment of cardiac muscle strips with 20 μM cTnT-C14 peptide had notable Ca^2+^-desensitization effects (decreased pCa50) at both short (2.0 μm) and long (2.3 μm) sarcomere lengths. On the other hand, the treatment with cTnI-C14 peptide did not decrease maximum force development of the cardiac muscle strips (Fig. 6A inset).

**Figure 6.**
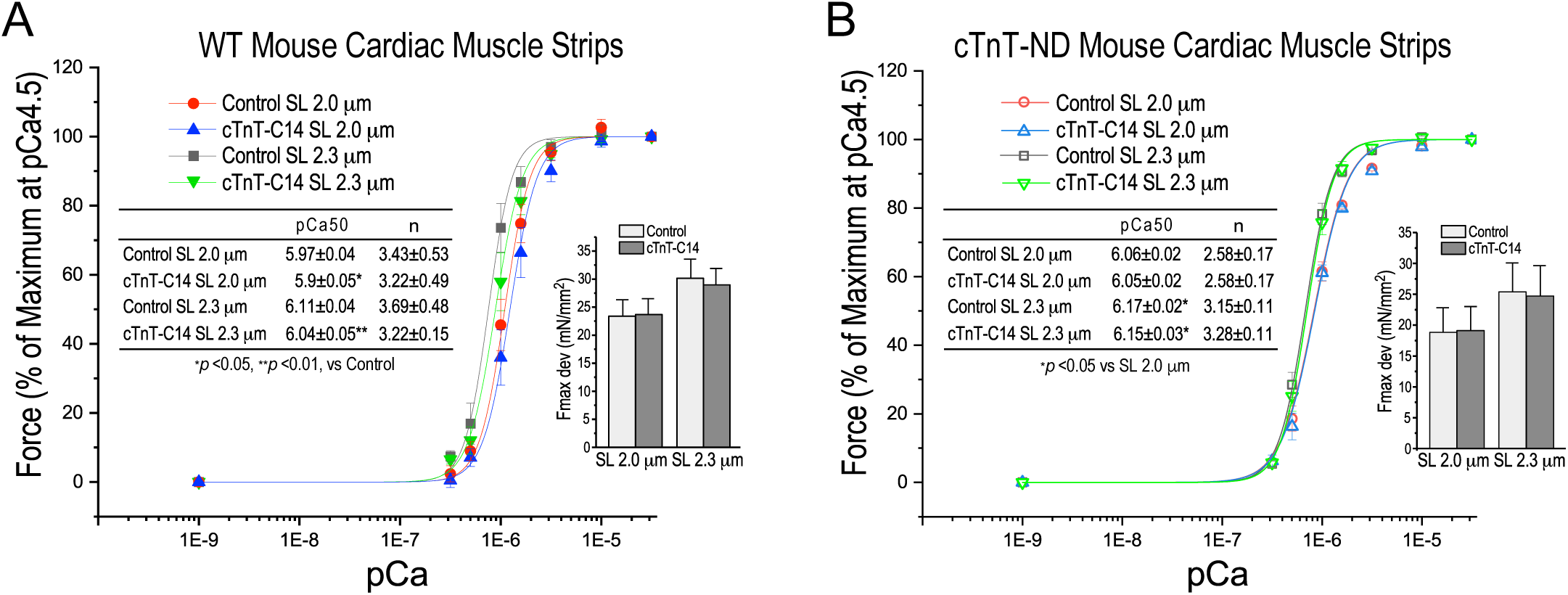
Effect of cTnT-C14 peptide on Ca^2+^-activation of permeabilized cardiac muscle strips. The force-pCa curves and inset tables and bar graphs show that (A) the addition of WT cTnT-C14 peptide to skinned strips of WT mouse left ventricular papillary muscle decreased Ca^2+^- sensitivity. At sarcomere lengths 2.0 μm and 2.3 μm, the treatment with 20 μM cTnT-C14 peptide had a notable Ca^2+^-desensitization effect on cardiac muscle strips (*P < 0.05, **P < 0.01 vs control at the same sarcomere length, SL), and (B) the adding of WT cTnT-C14 peptide to skinned strips of cTnT-ND mouse left ventricular papillary muscle in force-pCa measurements did not produce the Ca^2+^-desensitization effect while the expected positive response to increasing SL from 2.0 μm to 2.3 μm was retained (*P < 0.05). cTnT-C14 peptide did not decrease maximum force development (Fmax dev) of the cardiac muscle strips in both experiments. N = 5 fibers in each group.

The addition of WT cTnT-C14 peptide to skinned strips of cTnT-ND transgenic mouse left ventricular papillary muscle for Force-pCa studies did not no produce the Ca^2+^-desensitization effect (Fig. 6B). This finding suggests that the same mechanism had been utilized by the conformational effect of the restrictive deletion of the N-terminal variable region^19^ on reconfiguring the TnI-like inhibitory structure and function of the C-terminal end segment of cTnT-ND^7,8^. With the enhanced function of the endogenous Tm-binding site 3 in cTnT-ND in place^7,8^, the addition of exogenous cTnT-C14 peptide would not produce additive effect through the same mechanism. The similar and non-additive effects of the endogenous cTnT-ND-enhancement and the exogenous cTnT-C14 peptide on Ca^2+^-desensitization further support that the cTnT-C14 peptide retains its in situ physiological activity in adjusting cardiac muscle contractile kinetics. In addition to mimicking the function produced by cTnT-ND in vivo for a compensatory adaptation to myocardial contractility-afterload mismatch^25,19^, the physiologically restrictive functional effect of exogenous cTnT-C14 peptide is a plausible feature of a safe drug candidate.

Consistent with their diminished Tm- and actin thin filament-binding capacities, addition of the three HCM mutant cTnT-C14 peptides R278C, K280N or R286C, to skinned WT mouse left ventricular papillary muscle strips did not produce any notable functional effect (Fig. 7). The results demonstrate the loss of Tm- and thin filament-binding function as the underlying mechanism for these cardiac TnT point mutations to produce pathophysiological effects and cause myocardial over-activation in HCM.

**Figure 7.**
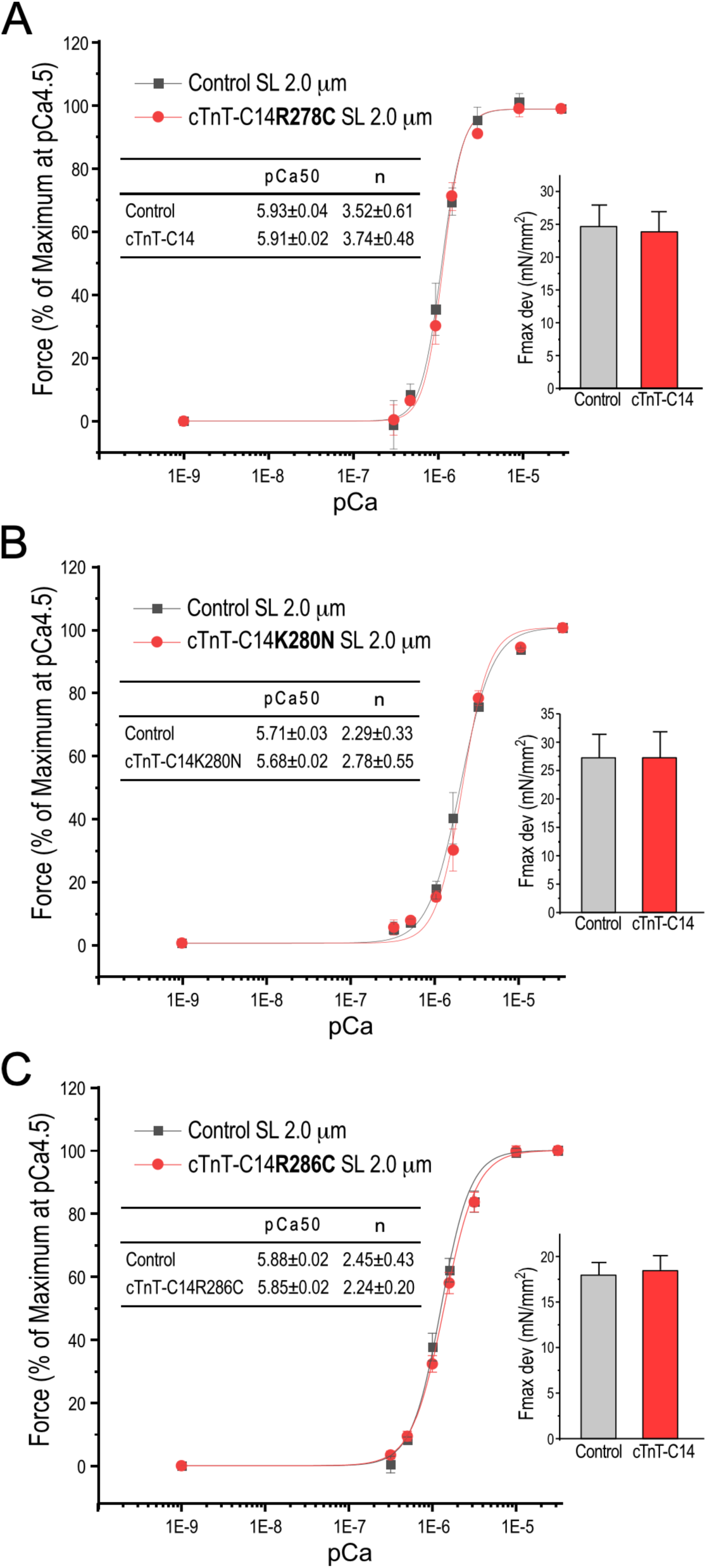
HCM mutations in cTnT-C14 peptide cause a loss of function in modulating myofilament Ca^2+^-sensitivity. Consistent with their diminished Tm- and thin filament-bindings, the force-pCa curves and inset tables and bar graphs show that the addition of 20 μM cTnT-C14 peptides containing R278C (A), K280N (B) or R286C (C) mutations to skinned strips of WT mouse left ventricular papillary muscle did not produce any detectable effect on Ca^2+^-activation of contraction, indicating a loss of function that underlies the pathophysiology of HCM caused by these cardiac TnT point mutations. The addition of mutant cTnT-C14 peptide did not decrease the maximum force development (Fmax dev) of the cardiac muscle strips. N = 4 strips for R278C, N = 5 strips for R286C and N = 3 strips for K280N studies.

## 4. Discussions

The present study investigated the functionality of the C-terminal end peptide of cardiac TnT, which contains the recently identified and characterized Tm-binding site 3 that possess a TnI-like inhibitory function in regulating contractile kinetics of cardiac muscle^7,8^. Using LSPR as an sensitive approach to measure protein-protein interactions and kinetics^26^, the studies of WT and three HCM mutants of human cTnT-C14 peptides for their interactions with Tm, Tm-F-actin thin filaments and F-actin alone solidified a mechanistic basis for physiological and pathophysiological studies. Followed by characterization of the functional effects of the WT and mutant cTnT-C14 peptides on the Ca^2+^-activation of cardiac muscle contraction, the present studies provides several findings that add new insights into the structure-function relationship of TnT in regulating striated muscle contraction and relaxation and demonstrate the loss function basis of point mutations in the C-terminal end segment of cardiac TnT in causing HCM. The functional characterization of the physiological functionality of the isolated cTnT-C14 peptide and its effects on adjusting the kinetics of cardiac muscle contractility are especially valuable for the development of a cardiac pumping efficiency-targeted treatment for heart failure.

### 4.1. The Tm-binding Site 3 in the C-terminal End Segment of Cardiac TnT Retains Physiological Functionality in the Form of Isolated Free Peptide

Pioneer work in the 1970-80’s by the Smillie group and others mapped the structure- biochemical function relationship of TnT using proteolytic and chemically cleaved fragments. The studies detected binding affinities for TnC, TnI, Tm and actin in the T2 fragment containing the C-terminal ∼100 amino acids^5^ and another Tm-binding site in the middle region cyanogen bromide fragment CB2^4^ (Fig. 1A). The precise locations of the TnC and TnI binding sites of TnT in the T2 region have been confirmed by high resolution crystallography studies of partial cardiac and skeletal muscle troponin complexes^27,28^. The crystallography structures of troponin complex did not resolve the middle region, the N-terminal variable region and the C-terminal end segment of TnT, likely reflecting their nature as flexible structural domains. Our follow up protein binding studies using molecular engineered TnT segments further mapped the Tm-binding site 1 that expands a large portion of the conserved middle region, and localized the Tm-binding site 2 to the beginning of the T2 fragment of TnT^6^. Revising the longstanding muscle thin model with two Tm-binding sites of TnT, we recently reported the identification and localization of a novel Tm-binding site 3 in the C-terminal end 14 amino acids segment of TnT^7,8^ (Fig. 1A).

To further establish the intrinsic function of the C-terminal end 14 amino acids segment of cardiac TnT, our present study focused on the form of free peptide in isolation from the TnT backbone. This strategy is based on a) this segment is a single exon-encoded genetic (and likely structural and functional) unit, b) it is a structure outside of the core of troponin complex as indicated by the crystallography data^27,28^ (Fig. 1A), and c) it is a uniquely highly conserved structure with a 100% conserved amino acid sequence in all mammalian cardiac TnT (Fig. 1B). Proving a hypothesis based on the above rationale that the C-terminal end 14 amino acids segment may retain functionality when isolated from the TnT core structure, our data show that free C-terminal end 14 amino acid peptide of human cardiac TnT retains the function of Tm- and thin filament-binding with physiologically relevant affinities (Figs. 3 and 4).

It is an important finding that the cTnT-C14 peptide also directly binds F-actin although in a non-saturable manner (Fig. 5). This activity is consistent with the previous reports that the T2 fragment of TnT (Fig. 1A) bound F-actin^24^. Our result now localizes the actin-binding activity of TnT to the C-terminal end segment, which is also a function retained by the cTnT-C14 peptide in isolation from the troponin backbone and apparently structurally integrated with the Tm-binding site 3. This finding lays a new platform for the structure-function model of troponin regulation of muscle contraction and relaxation.

### 4.2. Loss of Function Underlies the Pathological Impact of HCM Mutations in cTnT-C14 Segment

The C-terminal end segment of TnT encoded by the last exon of *TNNT2* gene is highly conserved in vertebrate species and has identical amino acid sequence in all mammalian cardiac TnT (Fig. 1B). This unique structural conservation reflects a high selection value during the functional evolution of mammalian hearts based on a stringent structure-function relationship. The loss of binding affinities of cTnT-C14 peptide for Tm, F-actin and Tm-F-actin thin filament caused by the HCM-associated mutations (Figs. 3, 4 and 5) confirms the functional importance of this regulatory structure and its stringent structural requirement.

It has been shown that R278C mutant of cardiac TnT increases Ca^2+^-sensitivity with reduced thin filament binding affinity^10,29^. The previous data are consistent with our finding in contractility studies that the three loss of function HCM mutations abolished the Ca^2+^- desensitization effect of cTnT-C14 peptides (Fig. 7). The loss of Tm-binding site 3 function in inhibitory regulation of cardiac muscle contraction is likely to hinder the coordinated transition from contraction to relaxation to cause impaired relaxation and hypercontractile phenotypes as shown by the study of Ala substitution for the positively charged Arg and Lys residues in the C-terminal end segment of TnT^9^.

As an allosteric regulatory structure, the C-terminal end segment of TnT should be sensitive to impaired conformational and functional dynamics. An interesting finding in our LSPR kinetic studies is that the three HCM mutations in the C-terminal end segment of cardiac TnT show a common feature of slowing down both association and dissociation of cTnT-C14 peptide in the interactions with Tm and Tm-F-actin filament (Figs. 3, 4 and 6). Fast kinetics of both association and dissociation rates of WT cTnT-C14 peptide’s interaction with Tm indicate a dynamic process where transitions occur readily between bound and unbound states during the rhythmic cycles of cardiac muscle contraction and relaxation, which is required for powering the heart function as an efficient pumping machine. While the association and dissociation rates are both impaired by the HCM mutations, the less hindered rate of association of the HCM mutant cTnT-C14 peptides (Tables 1-3) suggests that the mutations do not abolish the initial Tm-F-actin thin filament binding of the Tm-binding site 3 of cardiac TnT whereas the significantly slower thin filament dissociation rate caused by HCM the mutations may restrict the activation in the next cycle of cardiac muscle contraction. This observation suggests that the binding kinetics of the Tm-binding site 3 may be a critical factor in the rhythmic contractions of cardiac muscle and a sensitive target of pathologic mutations as well as therapeutic approaches.

### 4.3. Conformationally Modulated Functionality of the C-terminal End Regulatory Domain of TnT

An initial lead to our finding of the Tm-binding site 3 with regulatory function in the C-terminal end 14 amino acid segment of TnT^7,8^ was the observation that a restrictive deletion of the evolutionarily added N-terminal variable region of cardiac TnT in adaptation to inotropy- afterload mismatch^25^ restores an evolutionarily repressed TnI C-terminus-like^14^ Tm-binding inhibitory structure^17,18^. The restorability of an evolutionarily repressed molecular conformation and function in the C-terminal end segment of TnT is a physiological examples of the role of the N-terminal variable region in conformational modulation of TnT functions^30,31^. In this mechanism, the functional interactions of C-terminal end domain of TnT with Tm and F-actin are related to the overall conformation of TnT in the troponin complex.

It is worth noting that the molecular conformation of the TnI-like Tm-binding site of TnT can be modulated from a largely repressed state in intact cardiac TnT to the restored states in cTnT-ND and in a mini-TnT construct containing only the C-terminal portion of the T2 region^7,8^. In addition to the intramolecular conformational modulation mechanism, a previous study observed an effect of Ca^2+^-TnC on reducing the binding of the T2 fragment of TnT to F-actin^29^. These data let us propose that functionality of the C-terminal end segment of TnT may be dynamically changing between various allosteric states of troponin complex during Ca^2+^-regulated contraction and relaxation of cardiac muscle. In the meantime, similar to the C-terminal end 27 amino acid segment of cardiac TnI^17,18^ free cTnT-C14 peptide retains a functional structure in solution with Tm-binding activity (Fig. 3) and physiological function in modulating muscle contractility (Fig. 6). This retention of functionality in isolated cTnT-C14 peptide indicates that the activity of binding Tm and Tm-F-actin thin filament is an intrinsic function of the C-terminal end regulatory domain of TnT, which is in an unregulated active state when isolated from the TnT backbone with the overall conformational modulating effects removed.

### 4.4. Potential Use of cTnT-C14 Peptide as A Therapeutic Reagent

The present study demonstrates that the cTnT-C14 peptide can act at physiological concentration to adjust the Ca^2+^-sensitivity of cardiac muscle (Fig. 6). The appealing Ca^2+^- desensitization function of cTnT-C14 peptide presents an attractive value in therapeutic applications for the treatment of hypercontractile cardiomyopathies and diastolic heart failure. The nature of cTnT-C14 peptide with physiologically compatible dynamic interactions with Tm and Tm-F-actin filament provides a key benefit for pharmacological applications targeted at the modulation of the contractile kinetics of cardiac muscle to correct hypercontractility and hemodynamic efficiency of the rhythmic pumping cycles.

The finding that adding of cTnT-C14 peptide to cTnT-ND cardiac muscle strips did not produce additive effect on Ca^2+^-desensitization (Fig. 6B) indicates that the exogenous cTnT-C14 peptide utilizes the same myofilament regulatory mechanism as that of the C-terminal domain of endogenous cardiac TnT restored by conformational modulation of restrictive N-terminal truncation during a physiological adaptation in vivo^19,25^. This result is important by not only showing that the application of exogenous cTnT-C14 peptide safely utilizes an endogenous physiological mechanism but also demonstrating its function is a physiologically restraint effect on cardiac muscle contractility. This feature is highly plausible for a drug candidate for the treatment of diastolic heart failure by Ca^2+^-desensitization while avoiding over-desensitization.

In addition to adding insights into the mechanisms in the troponin regulation of striated muscle contraction and underscoring the pathophysiological basis of HCM mutations in the C-terminal end domain of cardiac TnT, the new findings in the present study highlight the cTnT-C14 peptide as a promising drug candidate for use as a pharmacological agent to modulate contractile function in the treatment of heart failure, especially diastolic failure.

## Acknowledgements

This study was supported by grants from the National Institutes of Health (HL127691 and HL138007 to J.-P.J).

## Novelty and Significance

The C-terminal end segment of TnT is a newly identified tropomyosin (Tm)-F-actin-binding structure with conserved amino acid sequence and a conformationally modulated TnI-like regulatory function. Mutations in this segment cause hypertrophic cardiomyopathy (HCM). The present study characterized the Tm- and F-actin-binding kinetics and myofilament Ca^2+^- desensitization activity of isolated C-terminal end 14 amino acids peptide of cardiac TnT (cTnT- C14). HCM mutations impair the peptide’s thin filament binding activity and diminish its physiological Ca^2+^-desensitization function. The data demonstrate the role of the cTnT-C14 segment in tuning contractile kinetics and the functionality retained in the form of free peptide establishes a potential therapeutic approach to adjusting cardiac muscle contractility to sustain the function of failing hearts.

